# Dopamine-induced Relaxation of Connectivity Diversifies Burst Patterns in Cultured Hippocampal Networks

**DOI:** 10.1101/2024.06.26.600923

**Authors:** Huu Hoang, Nobuyoshi Matsumoto, Miyuki Miyano, Yuji Ikegaya, Aurelio Cortese

## Abstract

The intricate interplay of neurotransmitters orchestrates a symphony of neural activity in the hippocampus, with dopamine emerging as a key conductor in this complex ensemble. Despite numerous studies uncovering the cellular mechanisms of dopamine, its influence on hippocampal neural networks remains elusive. Combining in vitro electrophysiological recordings of rat embryonic hippocampal neurons, pharmacological interventions, and computational analyses of spike trains, we found that dopamine induces a relaxation in network connectivity, characterised by a reduction in spike coherence. This relaxation expands the repertoire of burst dynamics within these hippocampal networks, a phenomenon notably absent under the administration of dopamine antagonists. Our study provides a thorough understanding of the roles of dopamine signalling in shaping functional networks of hippocampal neurons.

## Introduction

The brain operates as a dynamic and intricately interconnected network of neurons, where neurotransmitters choreograph the symphony of neural activity (Bullmore and Sporns, 2009; Friston 2011). Among these neurotransmitters, dopamine (DA) emerges as a central orchestrator, exerting profound effects across various brain regions (Wise, 2004; Schultz, 2007). Notably, the hippocampus stands out as a significant target of dopaminergic influence, receiving inputs from multiple brain nuclei (Swanson, 1982; Gasbarri et al., 1997; McNamara and Dupret, 2017; Takeuchi et al., 2016). The indispensable role of dopamine in hippocampal function is evident from previous research emphasising its involvement in spatial navigation (Xing et al. 2010; Kempadoo et al. 2016), memory formation (Shohamy and Adcock, 2010; da Silva et al. 2012) and even neuropsychiatric disorders such as schizophrenia (Grace, 2012; 2016).

*In vitro* cell cultures, combined with multi-electrode arrays (Kayama et al., 2018; Sasaki et al., 2019), offer compelling models for investigating neuronal activity in the hippocampus under the influence of dopamine. Previous studies have demonstrated that *in vitro* hippocampal neurons exhibit characteristic network-wide bursting activity (Suresh, 2016; Gonzalez-Sulser et al., 2012; Mody & Staley, 1994; Pimashkin et al., 2011). This activity can be experimentally manipulated by modulating synaptic transmission (Niedringhaus et al., 2013) or influencing dopamine-receptor sensitivities (Xie et al., 2011; Miyawaki et al., 2014; Li et al., 2017). Despite these insights into its cellular mechanisms (Yamada et al., 2002), there is a significant gap in understanding how dopamine impacts hippocampal network-level activity, which are critical for understanding the hippocampus’s role in learning, memory, and spatial navigation (Lisman, 2011; McNamara et al., 2014; Lisman et al., 2011). In the present study, we systematically explored the relationship between dopamine-induced alterations in network connectivity and the subsequent changes of burst patterns in primary hippocampal cultures. Employing a comprehensive approach involving electrophysiological recordings, pharmacological interventions, and computational analyses, we unveiled that dopamine induces a relaxation of connectivity within hippocampal networks, an effect characterised by a reduction in spike coherence in 1-ms bins. This relaxed state facilitates the emergence of a broader range of burst patterns within the neuronal ensemble, distinguished by clusters of rapid firing interspersed with periods of relative quiescence. In addition, we demonstrated that dopamine-induced changes in network connectivity correlate with the diversification of burst patterns, a correlation absent in samples treated with dopamine antagonists. These results suggest that dopamine plays a crucial role in shaping neural representations within the hippocampus, influencing information encoding, storage, and retrieval. Our study provides novel insights into the fundamental principles governing hippocampal information processing by dissecting the underlying mechanisms governing these phenomena.

## Results

We prepared a total of eight primary cultures from rat embryonic hippocampus and used MaxOne high-density microelectrode arrays (MaxWell Biosystems, Switzerland) to capture the spiking activity across several hundred active electrodes (Table 1). Initially, spikes were recorded in the absence of dopamine in the bath and were considered as the baseline condition. Subsequently, dopamine (DA) was gradually introduced into the bath, and spike recording sessions were conducted (Fig 1A). We systematically examined the effect of DA concentration on firing rate and found that dopamine concentrations ranging from 30 to 1000 µM consistently elevated the firing rate compared to baseline, in agreement with previous studies (Behr et al., 2000; Weiss et al., 2003). Consequently, in this study we analysed the effects of dopamine within this concentration range (see Supplementary Fig S1 and Methods for details). A linear mixed-effect model (see Methods for details) indicated that the neurons’ firing rate increased in response to the effects of dopamine (mean ± sem of the mean firing rate, 1.6 ± 0.3 Hz, 1.7 ± 0.3 Hz, 1.9 ± 0.4 Hz, 2.0 ± 0.3 Hz, 2.4 ± 0.3 Hz for baseline, 30, 100, 300, 1000 μM DA conditions, respectively; linear mixed-effects model, mean ± sem of DA coefficient, 0.24 ± 0.05, p < 0.00001). Examination of the raster plots of a representative sample revealed spontaneous, highly synchronised spikes, forming bursts within a 200-300 ms timeframe (Fig 1B). The presence of spike bursts, even evident under baseline conditions, implies the potential existence of neuronal networks influencing spiking dynamics within hippocampal cultures (Pimashkin et al., 2011; Suresh et al. 2016). In the following sections, we investigated the impact of dopamine on network connectivity and its subsequent effects on bursting patterns. Each analysis began with an examination of the effects on a representative sample, followed by group-level results and statistics.

**Figure 1:**
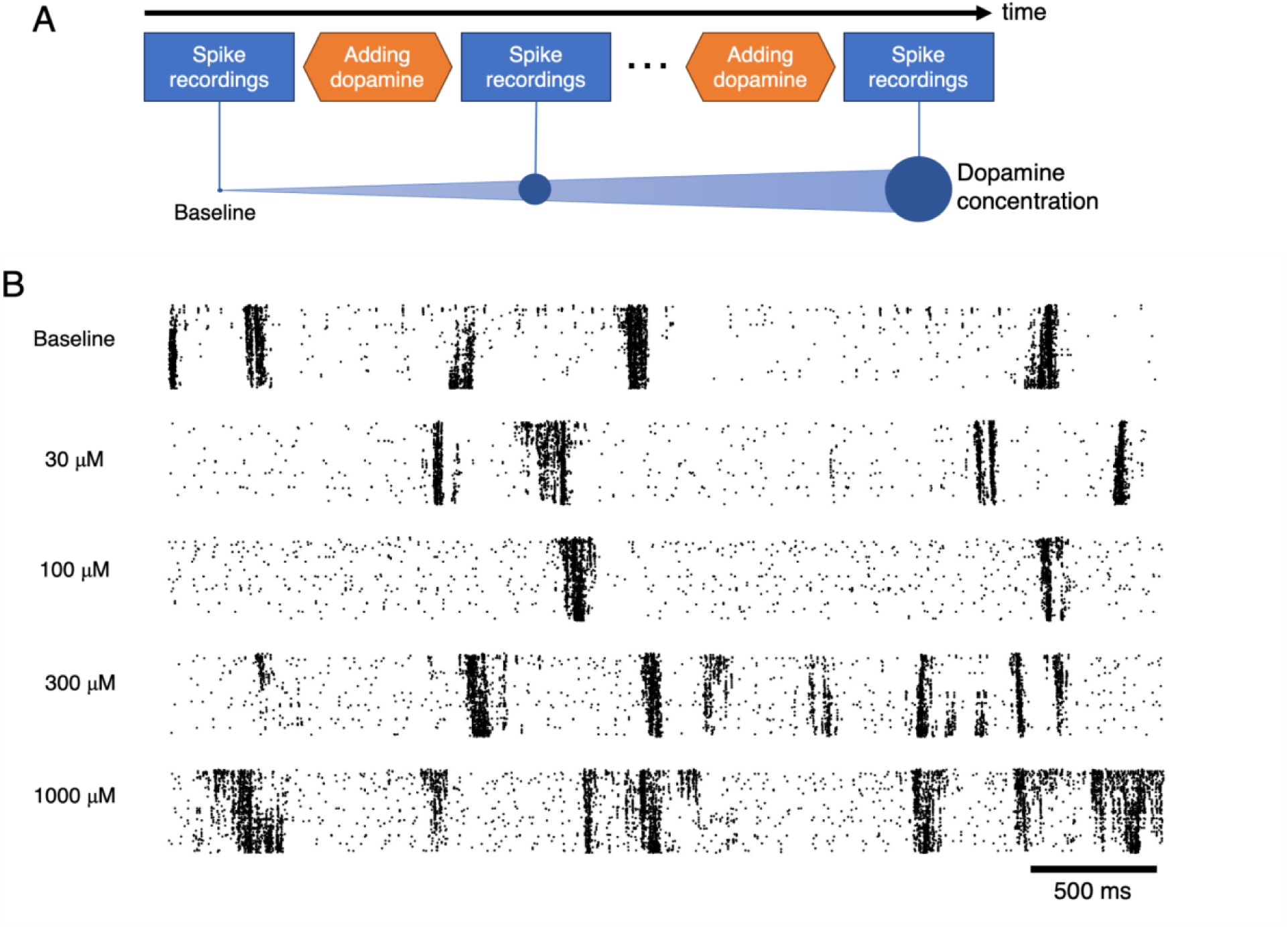
D**o**pamine **Experiment on Hippocampal Cultures Recorded by MaxOne Chip** A: The experimental scheme. B: Raster plots depicting spiking activity of active electrodes for the representative sample at baseline level (top) and at four levels of dopamine (DA) concentration (second to bottom, 30, 100, 300, 1000 μM). Note that we used this representative sample to illustrate subsequent analyses (see Figs 2-5).

**Table 1:** Summary of the data samples. Values in columns of # active electrodes and DIV indicate mean ± sem across samples. DIV – day in vitro.

| Sample group | # samples | # active electrodes | DIV |
| --- | --- | --- | --- |
| No-Antagonist | 8 | 768 ± 94 | 61.2 ± 38.7 |
| SCH23390 (10 μM) | 9 | 800 ± 30 | 57.8 ± 15.7 |
| Sulpiride (100 μM) | 12 | 756 ± 112 | 50.2 ± 11.3 |

### Dopamine reduces connectivity strength in hippocampal networks

To analyse the networks in hippocampal cultures, we constructed the connectivity matrix for each sample under a specific condition. A connectivity matrix represents functional connections between different electrodes derived from their spiking activity. Each row and column in the matrix correspond to individual electrodes, and the value indicates the connectivity strength, quantified as spike coherence in 1 msec bins (Fig 2A, see Methods for details). Comparing two conditions (baseline and 1000 μM, Fig 2B-C) in the representative sample revealed a notable reduction in connectivity strength between electrode pairs in the presence of DA (mean ± std of connectivity strength across electrode pairs, 0.04 ± 0.04 and 0.02 ± 0.03 for baseline and 1000 μM, respectively, Fig 2D). This reduction was consistent across all electrode pairs (Fig 2E), suggesting that dopamine broadly impacted network connectivity.

**Figure 2:**
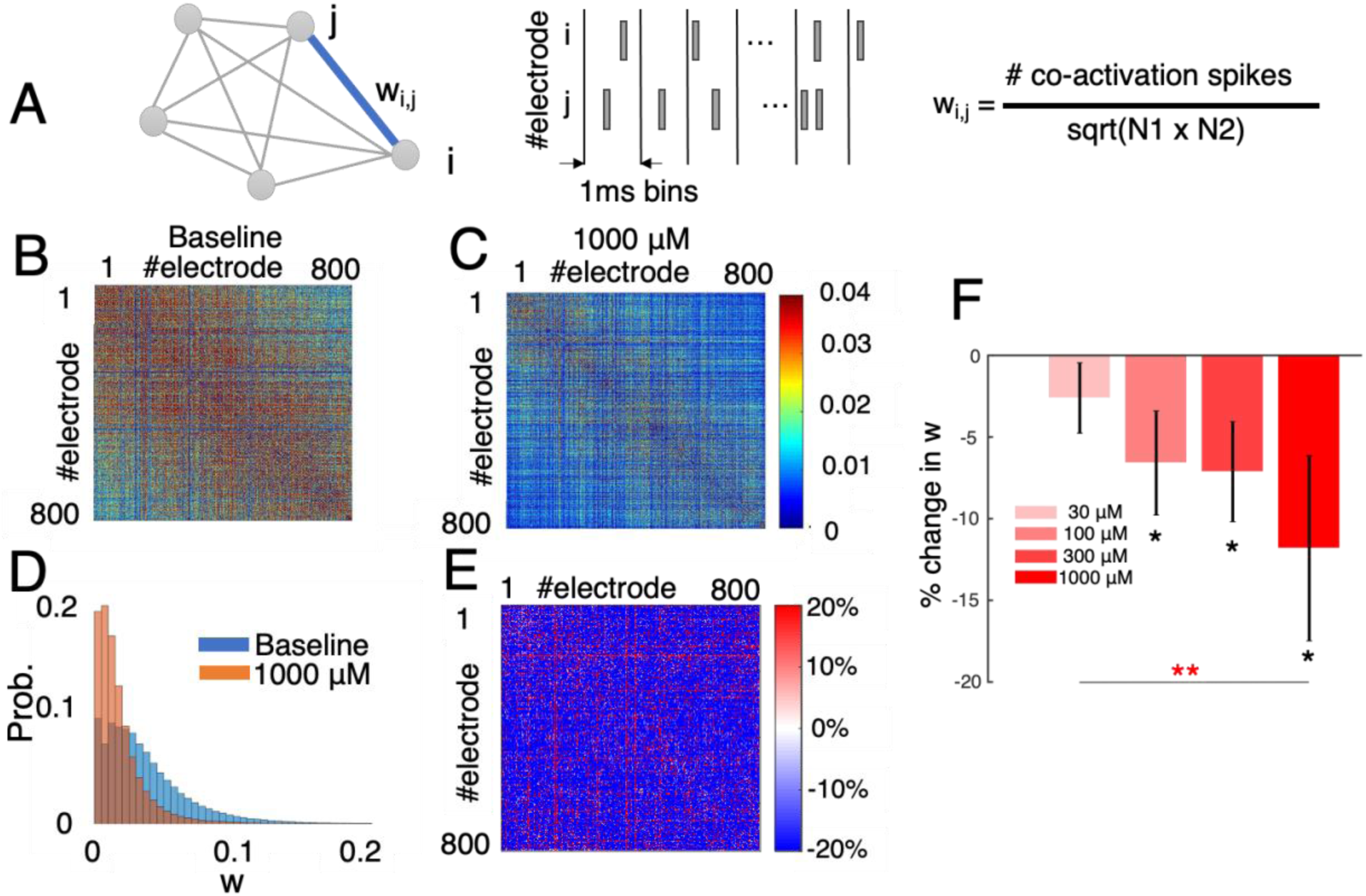
E**f**fects **of Dopamine on Connectivity Strength of Hippocampal Networks** A: Network analysis with the connectivity strength defined as co-activation of spikes within 1-msec bins. B-E: Connectivity matrix of the representative sample in the baseline condition (B) and when the DA concentration level reached 1000 μM (C), histograms of connectivity strength (D), and difference in connectivity matrix (E). F: Change in averaged connectivity strength, compared to the baseline level, for the four DA conditions (four shades of red, each for 30, 100, 300, 1000 μM) in n=8 samples. Bars with errors represent mean and s.e.m, respectively. Red asterisks indicate the significance level of the main effect of DA, computed with a linear mixed effects model. Black asterisks indicate the significance level of one-tailed t-tests against 0. * p < 0.05, ** p < 0.01.

Considering all samples, connectivity strength consistently diminished with increasing dopamine concentration levels. A linear mixed-effects model with the mean connectivity strength as the response variable captured this effect, showing a negative coefficient of DA concentration level (mean ± sem, –0.0013 ± 0.0004, p = 0.006). Post-hoc tests highlighted that, relative to baseline, the reduction in connectivity strength was statistically significant for all dopamine conditions except the 30 μM condition (mean ± sem of % change in the mean connectivity strength across samples: –3 ± 2%, –7 ± 3%, –7 ± 3%, –12% ± 6% for 30, 100, 300, 1000 μM DA conditions, respectively, Fig 2E). Importantly, this trend was observed irrespective of bin size (Supplementary Fig S3D, varying bin size from 1 to 50 msec).

### Dopamine increases the number of network modules and confines the connectivity within individual modules

In our study, we delved into the organisation of hippocampal networks in terms of distinct modules, each potentially responsible for specific aspects of computation or learning. To this end, we conducted modularity analyses of the connectivity matrix, aiming to identify a limited number of modules characterised by significantly stronger connectivity within modules compared to connections across modules (see Methods for details). Analysing the representative sample revealed three key observations when comparing the modules detected in the baseline condition with those under 1000 μM DA condition (Fig 3A-C). First, the modules were non-overlapping localised regions within the network (Fig 3A). Second, while two out of three modules in the baseline condition retained their structure, one module (depicted by purple dots in Fig 3A) was split into two distinct modules when exposed to 1000 μM DA concentration (indicated by purple and green dots in Fig 3B). Third, electrodes exhibited a higher likelihood of connectivity with other within-module electrodes under the 1000 μM DA condition compared to the baseline, as evidenced by the smaller number of across-module connections (black thick traces in Fig 3A-B). This was corroborated by the participation coefficient, a metric that quantifies the strength of connections between electrodes within their respective modules (see Methods for details), with higher coefficients indicating greater within-module connectivity (mean ± std of participation coefficient across electrodes, 0.55 ± 0.08, 0.62 ± 0.08 for Non and 1000 μM DA conditions, respectively, Fig 3C).

**Figure 3:**
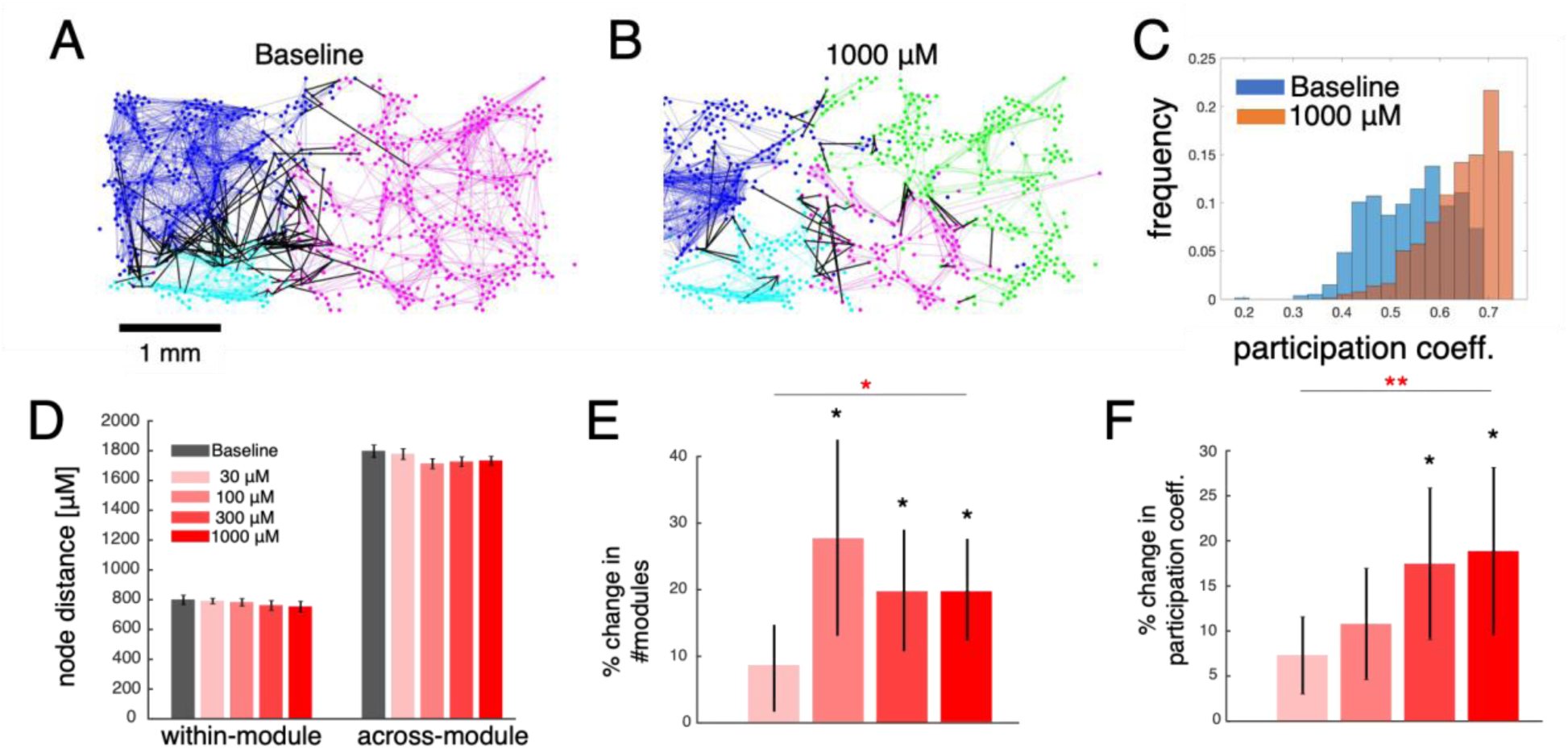
E**f**fects **of dopamine on network clusters and their connectivity** A-B: Spatial profiles of the presentative sample in the baseline condition (A) and 1000 μM condition (B). Each dot represents the relative position of a single electrode in the MaxOne high-density microelectrode array. Colours represent the modularity, where electrodes in the same module share the same colour. Lines connect the electrode pairs whose connectivity strength was greater than a threshold, defined by the top 1% in the baseline condition. Coloured thin lines connect electrodes in the same module and black thick lines connect electrodes in different modules. C: Histograms of participation coefficients of the electrodes in the baseline condition (blue bars) and 1000 μM condition (orange bars) for the representative sample. D: Distance of electrode pairs in the two pools of within-module and across-module for the baseline condition (grey bars) and the four DA conditions (red shaded bars). E-F: Changes in number of modules (E) and the mean participation coefficient (F), normalised by the baseline level, for the four DA conditions, n=8 samples. The convention was the same as in Fig 2. * p < 0.05, ** p < 0.01

All of these observations held true upon systematic examination across all samples. Specifically, the distance between electrodes within a module (∼800 um) was approximately half of that across modules (>1700 um), and these distances remained unchanged even under the influence of dopamine (Fig 3D), suggesting spatial confinement of the modules. Linear mixed-effects models revealed that elevated dopamine concentrations correlated with an increase in both the number of modules and participation coefficients (mean ± sem of DA coefficient across samples, 0.24 ± 0.10, p = 0.016 for number of modules and 0.03 ± 0.01, p = 0.0015 for participation coefficients). Relative to the baseline level, the number of modules exhibited a percentage change of 9 ± 7%, 27 ± 14%, 19 ± 8%, 19 ± 7% for 30, 100, 300, and 1000 μM DA conditions, respectively (Fig 3E). Similarly, participation coefficients displayed a percentage change of 7 ± 4%, 11 ± 6%, 17 ± 8%, 19 ± 9% for 30, 100, 300, 1000 μM DA conditions, respectively (Fig 3F). These increases were notably significant when dopamine concentrations ranged from 100 to 1000 μM.

### Dopamine diversifies activation patterns in burst events

We next sought to understand how dopamine would lead to alterations in the spike dynamics. To address this question, we focused on the activation patterns of bursts for two main reasons. Firstly, the networks exhibited bursting events characterised by high-frequency spiking involving a large number of electrodes (Fig 1B). Secondly, previous studies have indicated that the delays between the initial spikes within bursts of individual electrodes (activation patterns) reflect connectivity that remains relatively stable on the timescale of 100-200 ms (Raichman and Ben-Jacob, 2008; Pimashkin et al., 2011). Hence, we adopted a similar approach to Raichman and Ben-Jacob (2008) to quantitatively assess the changes in activation patterns across bursts induced by dopamine (see Methods for details).

Initially, bursts were detected by applying a threshold (0.2 standard deviations) to the trace of the number of active electrodes counted in 10-ms bins (Fig 4A). For each burst, the activation pattern was represented by a matrix *A*, where *A(i,j)* denotes the delays in milliseconds between the first spike of the *i*-th electrode and the first spike of the *j*-th electrode (Fig 4B). Our investigation indicated that the bursts endured for roughly 200 msec (mean ± sem, 204 ± 15 msec) with a firing rate of about 9 Hz (mean ± sem, 8.8 ± 0.4 Hz) and exhibited no notable change in either duration or burst firing rate under the influence of dopamine (data not shown). Note that the detected bursts remained consistent despite variations in the two parameters, threshold and bin size (Supplementary Fig S3A-B).

**Figure 4:**
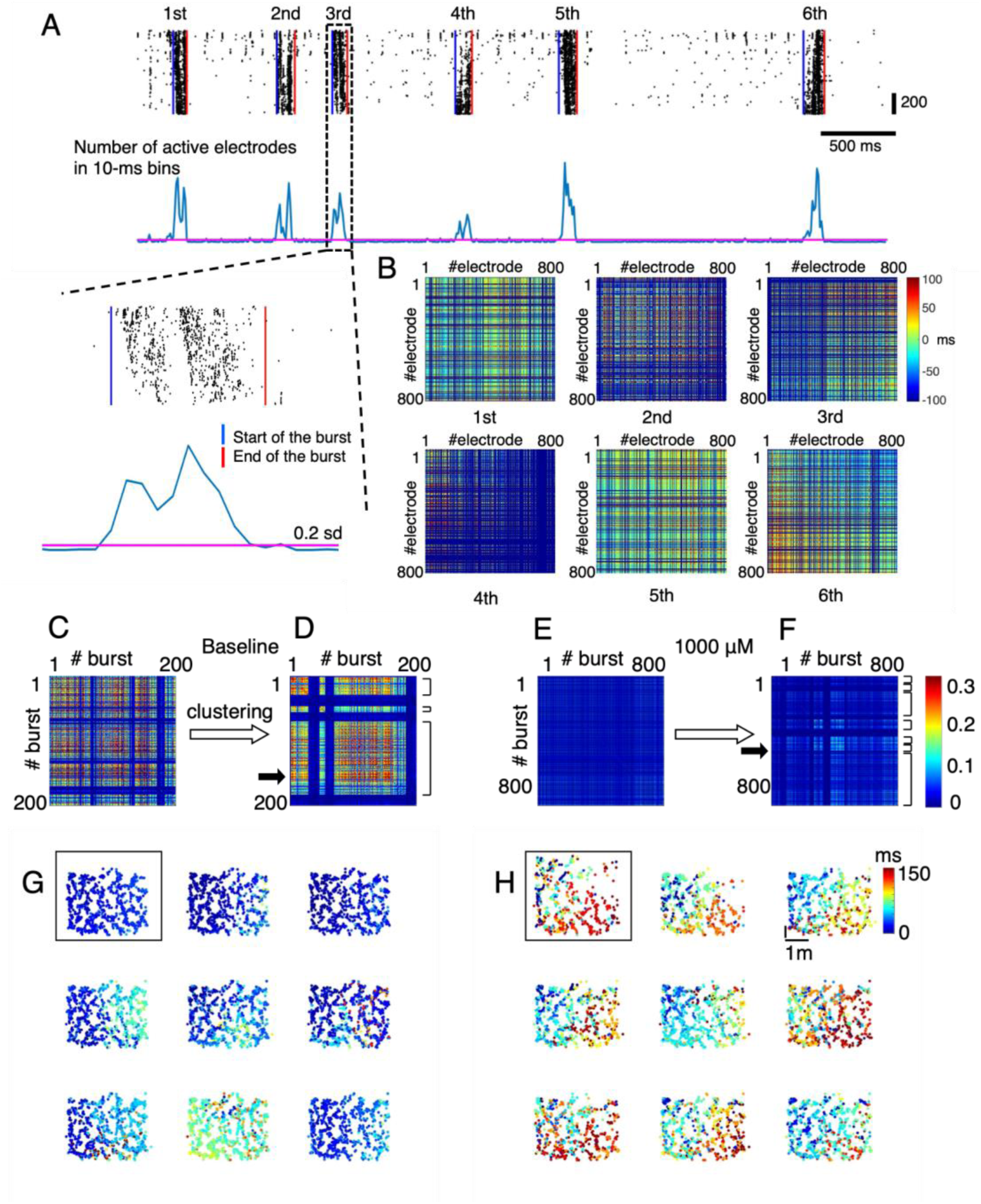
A**n**alysis **of Synchronized Bursting Events in a Representative Sample** A: Synchronised bursting events were identified based on timings when the number of active electrodes in 10-msec bins exceeded and fell below the threshold (magenta line), respectively. B: For each of the detected bursts, the activation pattern was constructed by capturing the differences in timing of the first spike in the burst between all electrode pairs. The six colour maps depict activation patterns of the bursts detected in Fig 4A. C-D: A similarity index matrix was constructed for all pairs of bursting events in the baseline condition of the representative sample (C). Using affinity propagation, we detected three activity motifs of bursting patterns, which are marked by the parentheses on the right side of the sorted similarity matrix (D). E-F: similar to C-D but for the condition of 1000 μM DA level. There were 7 activity motifs detected in this data condition. G: Scatter plots showing the relative position of the electrodes, with the colour, scaled from 0-150ms, illustrating the delay of the first spike timing of each electrode to the start of the burst. The scatter shown in the box was for a reference burst (indicated by solid black arrow in C), and the remaining 8 scatters represent the 8 bursts with the highest similarity to the reference burst. H: Similar to G but for the condition of 1000 μM DA level.

We then measured the similarity index between two bursts as the fraction of electrode pairs *(i,j)*, whose difference in *A(i,j)* of the two bursts is less than a temporal window of 50 ms. The similarity matrix was constructed by comparing the activation patterns between all detected bursts (Fig 4C). Next, we applied the affinity propagation algorithm on the similarity matrix to identify the clusters of burst patterns. We constrained our analysis on the clusters whose sizes are greater than one (we named them “activity motifs”) because of the following two reasons. First, in our dataset, 90% of the clusters detected by the affinity propagation contained only a single burst, but these single-burst clusters accounted for only about 20% of the total number of bursts (Supplementary Fig S4A-B). Second, the count of activity motifs was independent of the preference parameter in affinity propagation, which influences the number of clusters detected (Supplementary Fig S4C). Therefore, the activity motifs are burst patterns that occur frequently within the data, providing insights into underlying burst dynamics (see Methods for details). For the representative sample, we found that 1000 μM DA concentration increased the number of bursts 4-folds and the number of activity motifs 2-folds, and significantly reduced the similarity index in their activation patterns (mean ± std of similarity index, 0.11 ± 0.10, 0.02 ± 0.03 for baseline and 1000 μM condition, respectively, Fig 4C-F). Interestingly, the electrodes were co-activated within short temporal windows of <100 msec in the baseline condition (Fig 4G), but such activation was temporally and spatially divergent under the influence of dopamine (Fig 4H). Note that the change in bursting events was temporally irrelevant, both in the baseline and dopamine conditions (Supplementary Fig. S5).

Examination across eight samples confirmed that elevated DA concentrations increased the number of bursts (linear mixed-effects model with the number of bursts as response variable, mean ± sem of DA coefficient, 47.5 ± 15.4, p = 0.003; mean ± sem of % change in number of bursts relative to baseline level, 49 ± 43%, 70 ± 51%, 115 ± 83%, 161 ± 86% for 30, 100, 300, 1000 μM DA conditions, respectively, Fig 5A). In contrast, increased concentrations of dopamine led to a decrease in the mean similarity index between bursts. This was evidenced by a linear mixed-effects model with the mean similarity index as the response variable showing a negative coefficient of DA concentration level (mean ± sem, –0.013 ± 0.005, p = 0.019). More specifically, compared to baseline, the mean similarity index significantly reduced when the dopamine concentration reached 300-1000 μM (mean ± sem of % change in the mean similarity index: 0 ± 8%, –11 ± 10%, –24 ± 7%, –30 ± 12% for 30, 100, 300, 1000 μM DA conditions, respectively, as depicted in Fig 5B). It’s noteworthy that this reduction in similarity remained consistent irrespective of the size of the temporal window (Supplementary Fig S3C).

**Figure 5:**
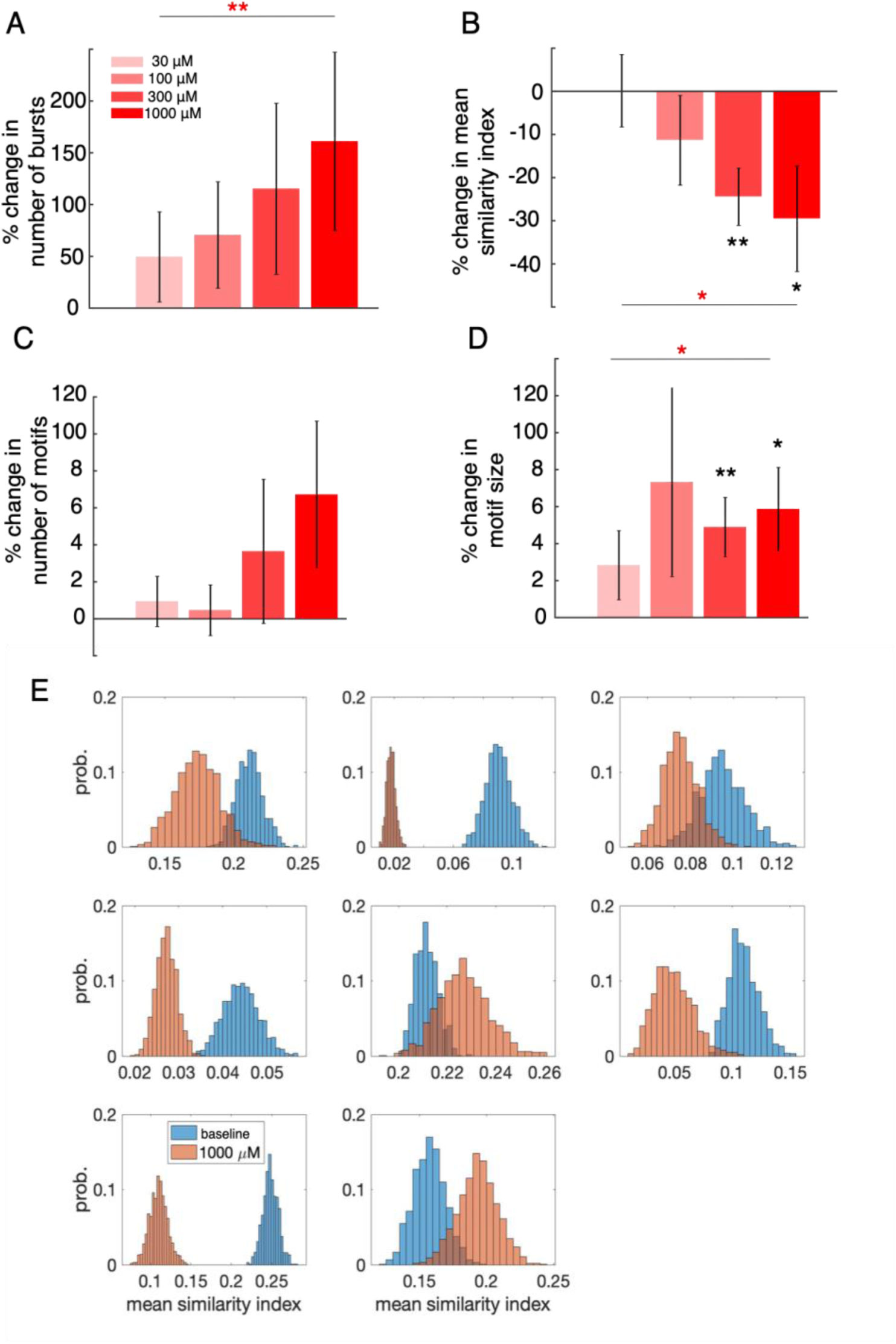
D**o**pamine **diversifies activation patterns in burst events.** A-D: Number of bursts (A), the mean similarity index between bursts (B), number of activity motifs (C), and the motif size (D), normalised by the baseline level, for the four DA conditions, n = 8 samples. Bars with errors represent mean and s.e.m, respectively. The convention was the same as in Fig 2. E: Probability distributions of the mean similarity index for the 8 samples following 1000 random resampling of the equivalent number of bursts in the baseline (blue bars) and 1000 μM DA condition (orange bars). * p < 0.05, ** p < 0.01

Compared to the baseline, the number of activity motifs increased by approximately 40-60% when the dopamine concentration reached 300-1000 μM. However, this increase was not statistically significant (linear mixed-effects model with the number of activity motifs as the response variable: mean ± sem of DA coefficient, 0.33 ± 0.23, p = 0.16, Fig. 5C). In contrast, higher dopamine concentrations significantly increased the motif size, which refers to the number of bursts in activity motifs. This was demonstrated by a linear mixed-effects model with motif size as the response variable (mean ± sem of DA coefficient, 6.65 ± 2.55, p = 0.013). The mean ± sem percentage change in motif size relative to the baseline level was 28 ± 19%, 73 ± 51%, 49 ± 16%, and 59 ± 23% for 30, 100, 300, and 1000 μM DA conditions, respectively (Fig. 5D).

One may ask whether the divergence of burst patterns was indeed influenced by elevated dopamine concentrations rather than an increase in the number of bursts (Fig 5A). To answer this question, for each sample, we randomly selected an equivalent number of bursts (n=100 or fewer if fewer bursts were present) in both the baseline and 1000 μM DA condition. We then computed the mean similarity index between the sampled bursts within the same condition. This resampling procedure was repeated 1000 times. As a result, six out of eight samples exhibited a significantly smaller mean similarity index in the 1000 μM DA condition compared to the baseline (Fig 5E). While the remaining two samples showed a slight increase in the mean similarity index in the 1000 μM DA condition, this trend was also evident in the original similarity index distribution (Supplementary Fig S6).

### Dopamine-induced decrease in connectivity strength causally decreases the similarity of activation patterns

In our final exploration, we investigated the causal relationship between the decrease in network connectivity and the resulting increase in burst pattern diversity through pharmacological intervention. Specifically, we tested the D2-receptor antagonist sulpiride and the D1-receptor antagonist SCH23390 (see Methods). We observed a notable effect of dopamine on connectivity strength (linear mixed-effects model, mean ± sem of DA coefficient, –0.0012 ± 0.0003, p < 0.001) but not the similarity index (p = 0.08) in n=12 samples treated with the D2-receptor antagonist sulpiride. Among the n=9 samples administered with the D1-receptor antagonist SCH23390, there were no significant dopamine effects identified (p > 0.2). Importantly, through an examination of inter-sample variability, we found a significant and positive relationship between the change in connectivity strength induced by dopamine and the change in similarity index across samples with no antagonist (linear regression, R^2^ = 0.22, p = 0.02, represented by red circles in Fig 6). Notably, this correlation was absent in samples treated with either SCH23390 (R^2^ = 0.04, p = 0.33, purple triangles) or sulpiride (R^2^ = 0.06, p = 0.19, grey squares). These findings suggest that the dopamine-induced decrease in connectivity strength within hippocampal networks directly contributes to the divergence in their activation patterns.

**Figure 6:**
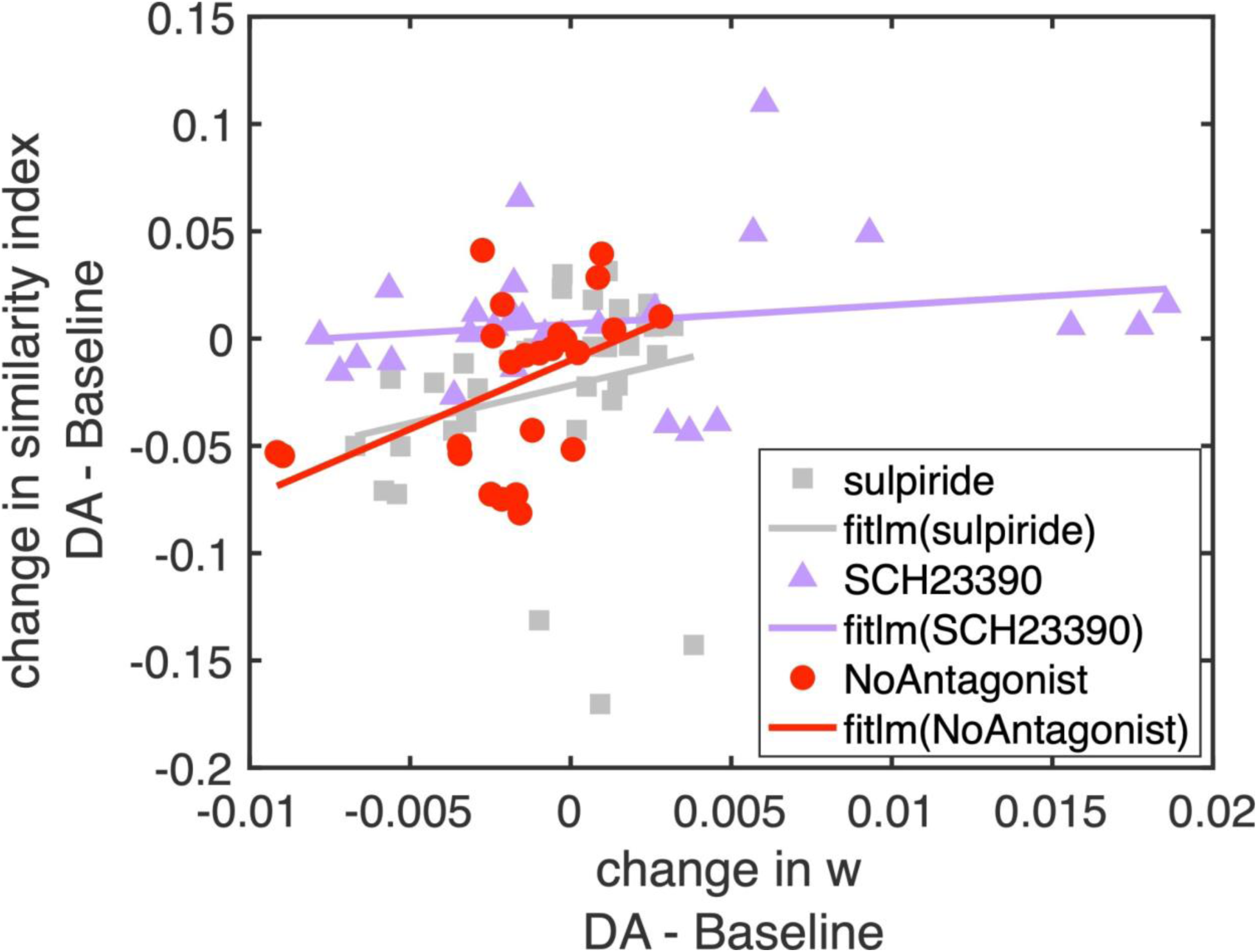
C**o**rrelation **of connectivity strength and similarity in activation patterns** The changes in similarity index (ordinate) were plotted against the changes in averaged connectivity strength (abscissa), and regression analysis was performed for three groups: samples with no dopamine antagonist (n=32, red circles), samples with sulpiride (n=48, grey squares), and samples with SCH23390 (n=35, purple triangles). Note that each sample has 4 DA concentration conditions in comparison with their baseline condition.

## Discussion

In this study, we examined the spiking activity of cultured hippocampal neurons under the influence of dopamine. By gradually administering dopamine, we observed a significant increase in the number of burst events, characterised by highly synchronised spikes across hundreds of electrodes within a 200-300 ms timeframe (Fig 1), consistent with previous findings (Li et al., 2017; Miyawaki et al, 2014). The high density of electrodes in our setup allowed us to monitor hippocampal neuron activity with cellular spatial and millisecond temporal resolution, enabling us to examine the effects of dopamine from a network-level perspective. Our observations revealed that dopamine administration resulted in a decrease in connectivity within cultured hippocampal networks, as demonstrated by a decrease in neural activity coherence (Fig 2). Additionally, dopamine increased the number of network modules, where each exhibited more confined connectivity (Fig 3). These findings imply that dopamine serves to restructure hippocampal networks, potentially influencing the flow of information processing in cognitive processes such as memory formation, decision-making, and attentional control (Lisman and Grace, 2005; Floresco and Grace, 2003; Schultz, 2007; Takahashi et al., 2008; Lemon et al. 2009; Smith and Greene, 2012). Supporting this hypothesis, our study also revealed that dopamine reduced the similarity in activation patterns between bursts and increased the number of activity motifs (Figs. 4-5). This expansion in the repertoire of burst dynamics may lead to a more distinct and separable encoding of information within the hippocampal circuitry (Sporns and Kötter, 2004; Chadwick and Hassabis, 2017; Ritchey et al., 2013; Staresina and Davachi, 2008). Interestingly, the diversification of firing patterns induced by dopamine was similarly reported by Miyawaki et al. (2014). Their study demonstrated that upon activation of dopamine receptors, different subsets of neurons were co-activated in sharp-wave events. This phenomenon was observed at a much higher spatio-temporal resolution in our study (Fig 4G-H). Most notably, we demonstrated that dopamine-induced decreases in connectivity correlate with reductions in the similarity index, a correlation absent in samples treated with either D1 or D2-receptor antagonist (Fig 6). An intriguing observation arises when considering that sulpiride, a medication used to treat schizophrenia, may potentially maintain the inherent structure of hippocampal networks while inhibiting the diversification of brain activity, a characteristic feature often observed in individuals with schizophrenia (Shiloh et al. 1997). Future studies of dopamine-mediated signalling using knock-in rats may provide valuable insights into the pathology of this disorder (Matsumoto et al., 2024).

It is important to clarify that we defined connectivity as spike coherence within 1-ms bins, rather than as functional connectivity (e.g., synaptic connectivity) between nodes (Cai et al., 2017; Kobayashi et al., 2019; Vareberg et al., 2024). This distinction is necessary because inferring functional connectivity from spike trains requires a relatively large number of spikes (Kobayashi et al., 2019), which is not feasible with our data. Our recordings have a mean firing rate of approximately 2 Hz over about 300 seconds per condition, resulting in an average of only 600 spikes per electrode. Consequently, this study focuses on the effects of dopamine on overall connectivity strength in the hippocampal network rather than on functional connectivity. It is also noteworthy that the dopamine concentration levels of 30-1000 µM examined in this study are much higher than those measured in an intact brain (Zetterström et al., 1983; Sharp et al., 1986; Borgkvist et al., 2012). However, similar DA concentrations have been reported as necessary to yield observable effects in the *in vitro* hippocampus (Behr et al., 2000; Weiss et al., 2003; Wang et al., 2020), a phenomenon also demonstrated in our investigation of firing rates (Supplementary Fig S1). Regarding the firing activity, hippocampal cultures typically display short-duration bursts, with significantly longer interburst intervals (as illustrated in Fig. 1B). Suresh et al. (2016) highlighted the crucial role of intrinsic membrane dynamics in burst initiation during interburst intervals, while synaptic processes operating on shorter timescales influence properties within-bursts and network connectivity. In our observations, the influence of dopamine on interburst activity remains uncertain, despite showing a modest increase in firing rate (linear mixed-effects models with firing rate in interburst intervals as response variable, p = 0.048). This suggests that dopamine may have a more profound impact on synaptic processes than on membrane dynamics. Nonetheless, further simulation work is required to dissect the specific effects of dopamine on spiking patterns at different timescales.

In summary, our study offers new insights into the causal relationship between dopamine-induced decreases in neural connectivity and the expansion of firing pattern repertoire within hippocampal networks. By elucidating the mechanisms underlying dopamine-mediated modulation of information processing, we contribute to a deeper understanding of the neural basis of hippocampus-dependent cognitive processes.

## Methods

### Ethical approval

Animal experiments were performed with the approval of the Animal Experiment Ethics Committee at the University of Tokyo (approval numbers: P29-3, P4-12) and according to the University of Tokyo guidelines for the care and use of laboratory animals. These experimental protocols were carried out in accordance with the Fundamental Guidelines for the Proper Conduct of Animal Experiments and Related Activities in Academic Research Institutions (Ministry of Education, Culture, Sports, Science and Technology, Notice No. 71 of 2006), the Standards for Breeding and Housing of and Pain Alleviation for Experimental Animals (Ministry of the Environment, Notice No. 88 of 2006) and the Guidelines on the Method of Animal Disposal (Prime Minister’s Office, Notice No. 40 of 1995). All efforts were made to minimise animal suffering.

### Animals

Embryonic day (E) 18 pregnant female Wistar/ST rats (Japan SLC, Shizuoka, Japan) were housed in a controlled environment with regulated temperature and humidity (22 ± 1 °C, 55 ± 5%) and maintained on a 12:12-hour light/dark cycle. Access to food and water was provided *ad libitum* throughout the duration of the study.

### Preparation

MaxOne high-density multi-electrode array (HD-MEA) chips (Maxwell Biosystems, Zurich, Switzerland) were pretreated and coated according to the manufacturer’s protocol. Briefly, 1% w/v Terg-a-zyme (Z273287, Merck, Darmstadt, Germany), dissolved in ultrapure water, was placed on each chip overnight for hydrophilization. On the next day, the Terg-a-zyme solution was removed. The chips and their lids were rinsed with ultrapure water three times and immersed in 70% ethanol for 30 min for sterilisation. After rinsing with ultrapure water and drying them, a coating solution was prepared, which was composed of 50% poly(ethyleneimine) (PEI) solution (P3143, Merck), 20× borate buffer (28341, Thermo Fisher Scientific, MA, USA), and ultrapure water at a final ratio of 140:4993:94867. The coating solution was sterilised by filtering and directly added to the HD-MEA. The chip was covered with its lid, incubated in 5% CO_2_ at 37℃ for 1 h, washed with ultra-pure water three times, and dried in air.

Primary cultures of hippocampal neurons were prepared from hippocampi of embryos of either sex as follows. The E18 rat was deeply anaesthetised by inhalation of an overdose of isoflurane, followed by intraperitoneal injection of a cocktail of xylazine (in saline, 12 mg/kg) and pentobarbital (in saline, 40 mg/kg). The rat’s uterus was carefully extracted and immersed in sterilised Ca^2+^– and Mg^2+^-free Hanks’ balanced salt solution (HBSS(-)) on ice, after which the mother rat was euthanized. The embryonic brains were extracted under microscopy and immersed in HBSS(-). The hippocampi were then dissected out. For the next step, Geltrex (A1569601, Thermo Fisher Scientific) was added to the coated HD-MEA chip, after which it was incubated in 5% CO_2_ at 37℃ for 1 h.

After dissection, the hippocampi were minced using a surgical knife, collected in additional HBSS(-), treated with 0.05% Trypsin-EDTA (25300062, Thermo Fisher Scientific), and incubated at 37°C for 15 min, followed by further incubation with 1% DNase I (in saline) (10104159001, Merck) at room temperature for 5 min and removal of the supernatant. Tissues were washed in HBSS(-) and incubated (in a bath) at 37°C for 5 min, after which the supernatant was removed. This procedure was repeated three times. Tissues were then triturated using a fire-polished Pasteur pipette in Neurobasal medium (12349015, Thermo Fisher Scientific) containing 2% B27 supplement (17504044, Thermo Fisher Scientific), 0.5 mM glutamine, 25 μM glutamate, 1 mM 4-(2-hydroxyethyl)-1-piperazineethanesulfonic acid (HEPES), 0.5% penicillin-streptomycin (09367-34, Nacalai Tesque, Kyoto, Japan), and 10% horse serum (26050088, Thermo Fisher Scientific), which we called plating medium. The tissues were further filtered through 100-μm-pore cell strainers, and centrifuged at 700 × g at 25°C for 5 min. The supernatant was replaced with a plating medium, which produced suspension. The suspension was mixed with nigrosin (198285, Merck) at a ratio of 1:1 to count the cells and calculate cell density on a hemocytometer (erythrocytometer). The initial cell concentration on Day 0 *in vitro* (DIV 0) was adjusted to 6 × 10^3^ cells/μl unless otherwise specified; the initial cell density was ∼6 × 10^3^ cells/mm^2^ because the whole area on the chip and cell drop volume were ∼50 mm^2^ and 50 μl, respectively. Cells were dispersed and seeded on the HD-MEA and incubated in 5% CO_2_ at 37°C for 1 h under maintained humidity, after which the plating medium was added to the MEA.

The feeding medium consisted of 2% B27, 1 mM HEPES, 0.5 mM glutamate, and 0.5% penicillin-streptomycin in Neurobasal medium and was used when the medium was replaced except for DIV 2. On DIV 1, the full medium was replaced with the feeding medium. On DIV 2, the full medium was replaced with the feeding medium including 5 μM cytosine arabinoside (AraC) to suppress the non-neuronal cell proliferation. On DIV 3, the full medium was refreshed. Afterwards, half of the medium was replaced twice a week. Every time the medium was replaced, the condition of the cultured neurons was checked with the eye to confirm that there was no evidence that active neurons were obviously contaminated, damaged, or dying.

### *In vitro* electrophysiology

The MaxOne recording system (Maxwell Biosystems) including the CMOS-based HD-MEA and MaxLab Live software (Maxwell Biosystems) including predefined pipelines were used to record and visualise the extracellular action potentials of the cultured hippocampal neurons. The HD-MEA chip had 26,400 platinum electrodes, organised in 220 × 120 pixels. The active sensing area was 3.85 × 2.10 mm^2^. The centre-to-centre electrode distance was 17.5 µm. Neural activity was measured at a sampling frequency of 20 kHz simultaneously from up to 1,024 channels in experimenter-based configurations with 10-bit analog-to-digital converter resolution.

The recording system was situated inside a 5% CO2 incubator at 37 ℃. Spontaneous neuronal activity was recorded from all channels using the ‘Activity scan’ module for 10 minutes, extracting the most ‘active’ 1,024 channels based on the firing rate. Identification of active channels subsequently allowed us to monitor collective neuronal activity using the ‘Network’ module for 5 minutes. The ‘Network’ recording was performed as follows. First, (1) dopamine (50 μM, 1 μl) was added to the chip (containing 500 μl medium) to achieve a final concentration of 0.1 μM dopamine. Neural activity was recorded at 0.1 μM dopamine using the ‘Network’ module. Next, dopamine was added in varying amounts ((2) 50 μM, 2 μl; (3) 50 μM, 7 μl; (4) 500 μM, 2 μl; (5) 500 μM, 7 μl; (6) 5 mM, 2 μl; (7) 5 mM, 7 μl; (8) 50 mM, 2 μl; (9) 50 mM, 7 μl; (10) 500 mM, 2 μl; (11) 500 mM, 7 μl; (12) 1 M, 10 μl) and neural activity was recorded at the following concentrations: (2) 0.3 μM; (3) 1 μM; (4) 3 μM; (5) 10 μM; (6) 30 μM; (7) 100 μM; (8) 300 μM; (9) 1 mM; (10) 3 mM; (11) 10 mM; (12) 30 mM. We systematically examined the firing rate of the samples at all recorded concentration levels and found that concentrations ranging from 30 µM to 1 mM consistently elevated the firing rate compared to baseline (Supplementary Fig S1). When the concentration level exceeded 1 mM (i.e., conditions (10)-(12)), the electrodes became completely silent (data not shown). Therefore, we focused on analyzing the effects of dopamine within the 30 µM to 1 mM concentration range. In antagonist experiments, sulpiride (25 mM, 2 μL) or SCH23390 (2.5 mM, 2 μL) was added to the chip to achieve a final concentration of 100 μM or 10 μM, respectively, before recording session (1). Recordings were regularly performed once a week.

### Preprocessing of *in vitro* spike trains

A considerable number of electrodes had relatively low activity during recordings. To address this issue, we implemented the following procedure for electrode selection. Initially, we assessed the number of active electrodes in time bins of 10 msec using all available electrodes. Subsequently, we systematically identified the top 90% of active electrodes based on their firing rates and determined the number of active electrodes using this subset. We then computed the correlation in the number of active electrodes between the two conditions. This process was repeated for subsets ranging from the top 80% to the top 10% of active electrodes. We found that the correlation remained consistently high (>0.99), even when considering only the top 50% of active electrodes (refer to Supplementary Fig S2). Based on these findings, we established that the correlation remained robust irrespective of the proportion of active electrodes considered. Consequently, in our analyses of all the samples, we opted to use the top 80% of active electrodes while discarding the remaining 20% of electrodes. This approach ensured a standardised and representative selection of electrodes for our analyses, enhancing the reliability and consistency of our results.

### Network analysis

In our study, networks were represented by connectivity matrices, where rows and columns corresponded to electrodes. The entries in these matrices indicated connectivity strength, determined by the number of spikes co-activated within 1 msec bins:

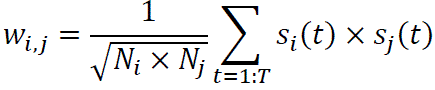

Here, *s_i_(t)* represents the number of spikes from the *i*-th electrode in the *t*-th bin, and *N_i_* represents the total number of spikes from the *i*-th electrode over the entire recording time *T*.

The modularization of the network involved dividing it into non-overlapping groups of electrodes, aiming to maximise the number of within-group weights while minimising the number of between-group weights. To detect these modules, we used the Brain Connectivity Toolbox (Rubinov and Sporns, 2010), employing the method of optimal modularity (Newman, 2006). If a module consists of only one electrode, it is reassigned to the nearest module based on its position.

Additionally, we computed the participation coefficient for each electrode. This coefficient measures the strength of a node’s connections within its module (Guimerà and Nunes Amaral 2005). The distance between nodes was defined by the Euclidean distance of the relative positions of the electrodes on the MaxOne chip.

### Detection of bursting events and evaluation of activation patterns in bursts

To detect bursting events, spikes were first binned into 10-msec intervals. The onset and offset of each burst were then identified by determining timings when the number of active electrodes exceeded and subsequently fell below a predefined threshold. This threshold was consistently set at 0.2 standard deviations. Bursts lasting less than 50 msec were excluded.

We adopted a similar approach to Raichman and Ben-Jacob (2008) to quantitatively assess the activation patterns across bursts. For each burst, the activation pattern was characterised by a matrix *A_k_*, where *A_k_(i,j)* represents the delay in milliseconds between the first spike of the *i*-th electrode 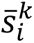 and the first spike of the *j*-th electrode 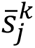 in the *k*-th burst:

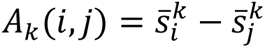

If an electrode did not fire during a particular burst, its value was set as NULL.

The similarity index *S(A_p_, A_q_)* between burst *p* and *q* was computed as follows:

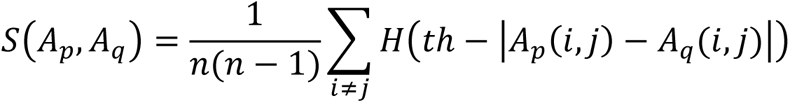

Where *n* denotes the number of electrodes, H is the Heaviside step function, defined as *H(x) = 0* if *x < 0* and *H(x) = 1* if *x ≥ 0*, and *th* is a time-threshold parameter. In this equation, *S(A_p_, A_q_)* represents the fraction of neuron pairs *(i,j)* for which the difference in delays between the two bursts is less than *th*: |*A*_*p*_(*i*, *j*) − *A*_*q*_(*i*, *j*)| < *t*ℎ. The summation is conducted only over neuron pairs *(i,j)* that did not receive NULL values in either burst (i.e., both neurons fired a spike in both bursts). In our analysis, we set *th* = 50 msec to reflect the average spike precision in bursts. We also tested other values of *th* ranging from 10 to 100 msec, and the results remained consistent (Supplementary Fig S2).

### Detection of repeating motifs from the similarity matrix

We applied the affinity propagation algorithm (Frey and Dueck, 2007) on the similarity matrix to identify clusters of burst patterns. Affinity propagation identifies exemplar points within the dataset and forms clusters around these exemplars. In our analysis, the algorithm used the similarities between pairs of burst patterns (described above), then iteratively updated responsibility and availability messages to determine which points would serve as exemplars. The responsibility message quantified how appropriate it was for each point to be chosen as an exemplar, while the availability message indicated how suitable it was for a point to be assigned to a particular exemplar. Through these iterative updates, the algorithm resulted in clusters where each burst pattern was assigned to the most suitable exemplar. The affinity propagation algorithm automatically determines the optimal number of clusters based on the preference parameter *p*(*i*), which indicates the preference that the *i*-th data point is chosen as a cluster exemplar. We set all preference values to the median of the similarity matrix. We defined activity motifs as clusters containing more than one burst and motif size as the number of bursts in each activity motif.

### Statistical analysis

We used linear mixed-effect models to statistically evaluate the influence of dopamine on the metrics of interest, such as connectivity strength and similarity index. Each model was structured as *Metric* ∼ *Concentration Level* + *(1 | Sample)*, where *Metric* is the metric of interest, *Concentration Level* represents the 10-base logarithm of dopamine concentration in the bath (with a value of 0 for the baseline condition), and *(1 | Sample)* accounts for repeated measures within each sample, thus adding a random effect for the intercept. Note that the logarithm represents the quantity of dopamine added during the experiments and does not alter the observed trend in the data. This analysis was conducted using the MATLAB function *fitlme*, from which we obtained the coefficient of *Concentration Level* and its corresponding significance p-value. Additionally, to assess the significance of dopamine-induced changes in these metrics relative to the baseline, we applied the following normalisation: *(Metric_DA_ – Metric_Baseline_) / Metric_Baseline_*, where *Metric_Baseline_* denotes the corresponding metric in the absence of dopamine. Subsequently, one-tailed t-tests were employed to determine whether the distributions of the metrics were significantly different from zero, representing the baseline level.

To evaluate inter-sample variability depicted in Fig 6, we employed a linear model to analyse the relationship between the amount of changes in connectivity strength and the amount of changes in the similarity index across samples. This analysis was conducted using the MATLAB function *fitlm*, from which we obtained the coefficient of determination (R^2^) and significance p-value. Throughout the paper, the significance levels were determined as follows: n.s (not significant) for p > 0.05, * for p < 0.05, ** for p < 0.01. These significance levels were used to interpret the statistical significance of the observed relationships. Unless specifically stated elsewhere, the mean ± sem of the statistics across samples were reported.

## Authors contributions

YI, NM and AC designed the study. NM and MM performed the experiments. HH and AC analysed the spiking data. All authors discussed the results and contributed to the final manuscript.

## Data and code availability

The customised MATLAB code of the analyses and spiking data of the samples was hosted publicly on github, accessible via https://github.com/hoang-atr/MaxOne.

## Acknowledgements

This study was supported by JST ERATO (JPMJER1801, “Brain-AI hybrid”). HH and AC were supported by Grants-in-Aid for Transformative Research Areas (22H05160). HH were partially supported by Grant Number JP23dm0307002, Japan Agency for Medical Research and Development (AMED).

## Supplemental Information

**Supplementary Figure S1:**
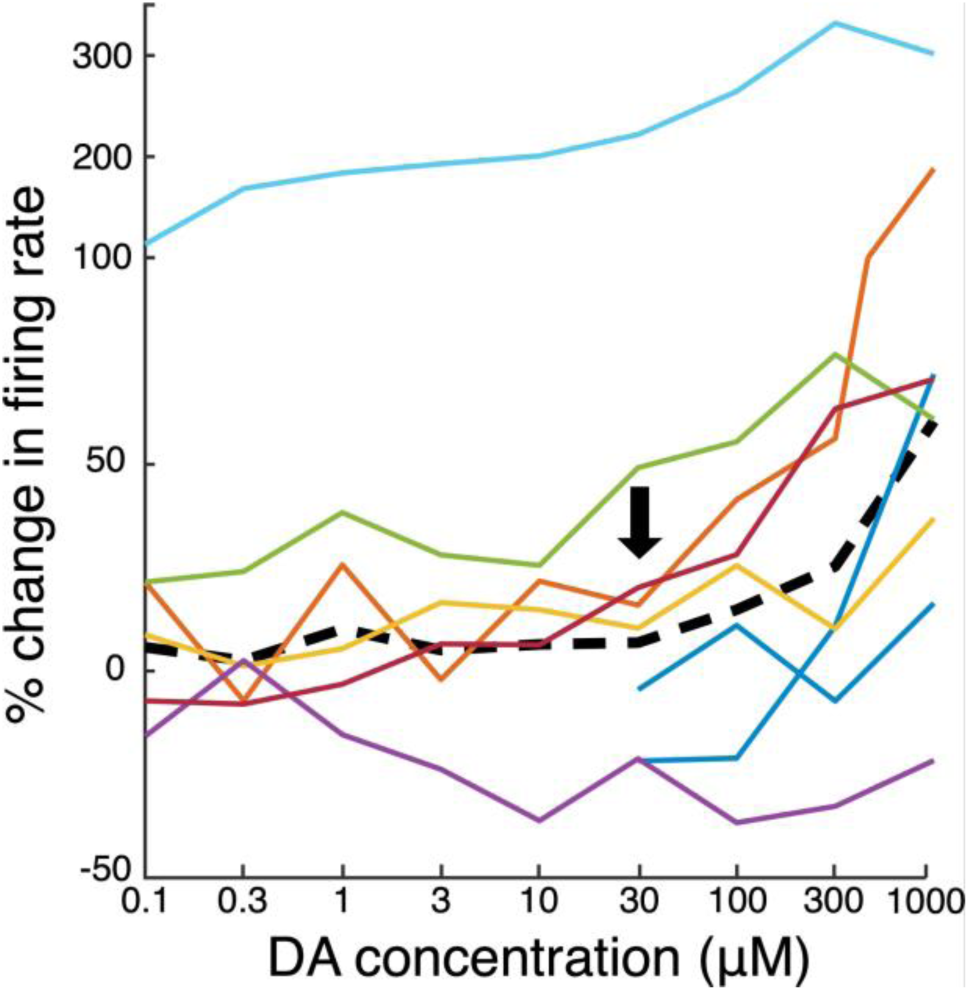
Change in firing rate compared to baseline for 8 samples (colored lines) by DA concentration ranging from 0.1 to 1000 µM. Note the scale of the ordinate to highlight an outlier sample (light blue line). The thick dashed line indicates the mean across samples excluding the outlier. The firing rate began to increase when the DA concentration reached 30 µM (black arrow). Additionally, note that two samples have missing data for the 0.1 to 10 µM conditions (blue lines).

**Supplementary Figure S2:**
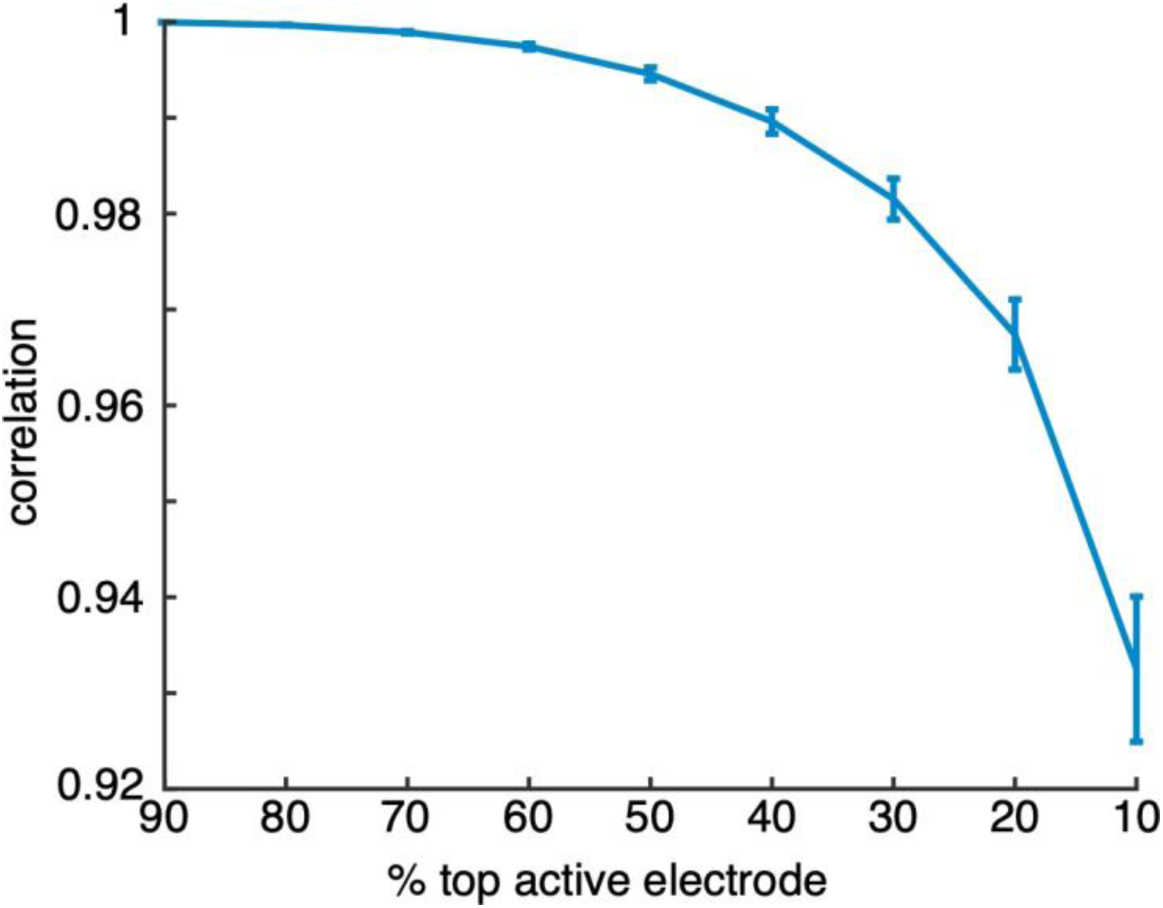
Selection of active electrodes by comparing the trace of number of active electrodes in 10 msec bins between the condition of all the available electrodes are selected and the condition of the top 90-10% of active electrodes are selected. The correlation was averaged for n=8 No-Antagonist samples with error bars indicating sem.

**Supplementary Figure S3:**
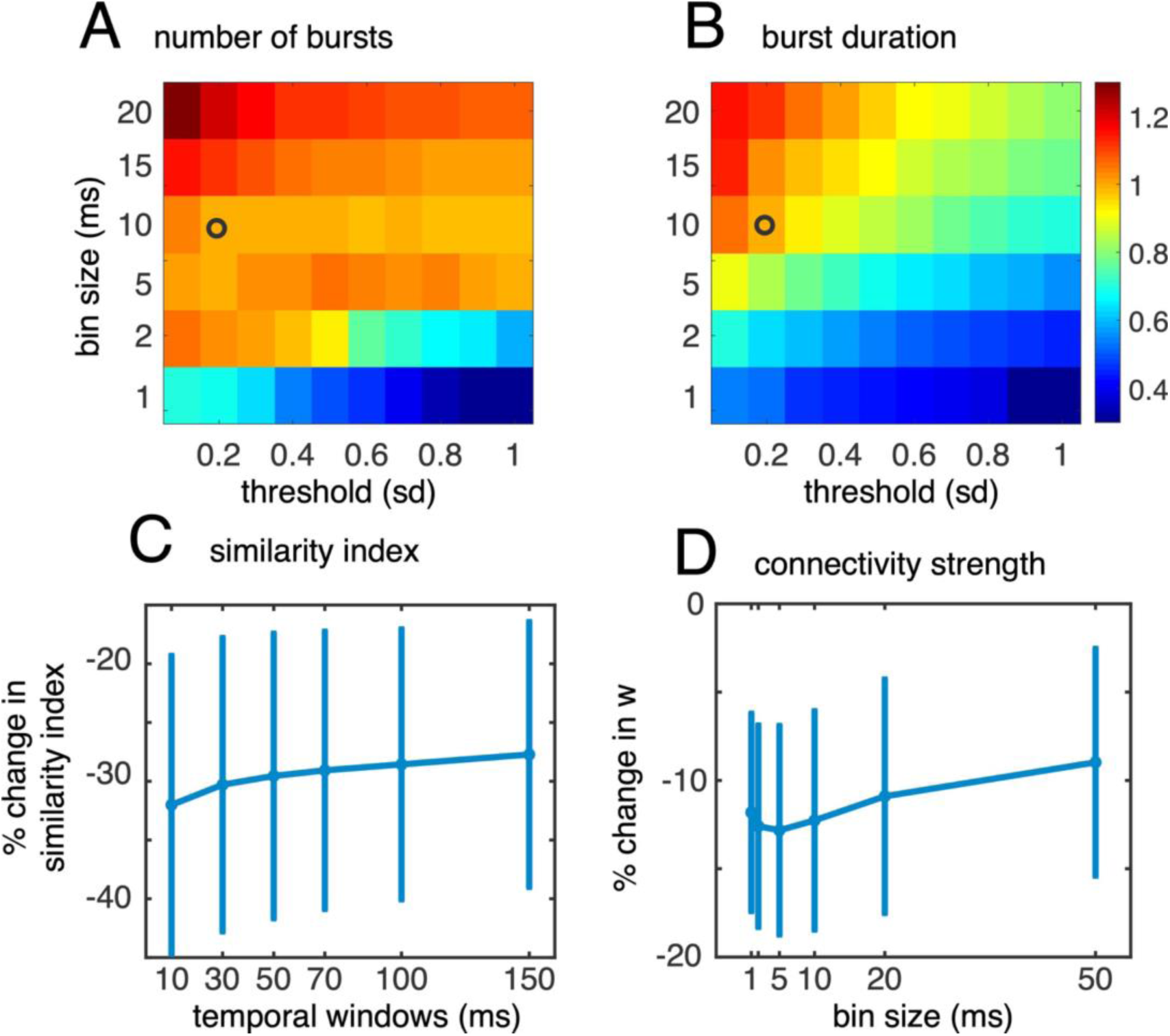
Validation of the hyper-parameters. A-B: Heatmaps depicted the average number of bursts (A) and the average burst duration (B) identified across 8 samples under baseline conditions, while systematically altering two parameters: the bin size of active electrodes and the threshold. These values were standardised against those obtained using the parameters detailed in the main text (bin size of 10 msec and threshold of 0.2 standard deviations, denoted by black circles). C-D: The decrease in the mean similarity index (C) and mean connectivity strength (D) among the 8 samples under 1000 μM DA condition compared to the baseline level was assessed across various parameters of temporal window for similarity index (C) and bin size for spike coherence (D). Line with error bars indicated the mean and sem across n=8 samples.

**Supplementary Figure S4:**
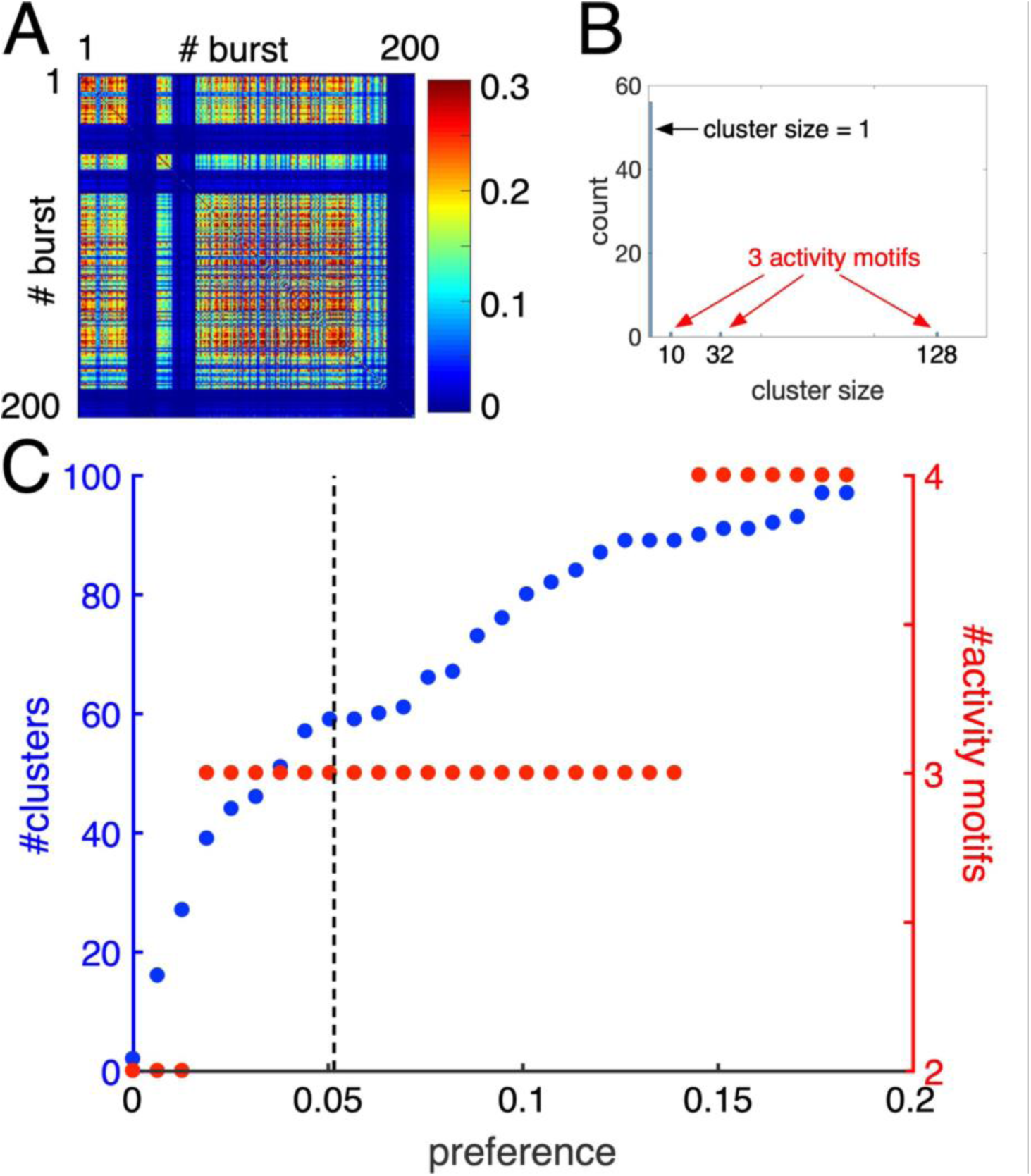
Detection of activity motifs via affinity propagation clustering for a representative sample. A: The similarity matrix of bursts in the baseline condition for the representative sample, sorted by clusters identified by the affinity propagation algorithm (corresponding to Fig 4D in the main text). B: Out of the 59 clusters detected, only three clusters with sizes greater than one were classified as activity motifs. C: The relationship between the number of clusters and the number of activity motifs as a function of the preference parameter of affinity propagation. The dashed line represents the median value of the similarity matrix.

**Supplementary Figure S5:**
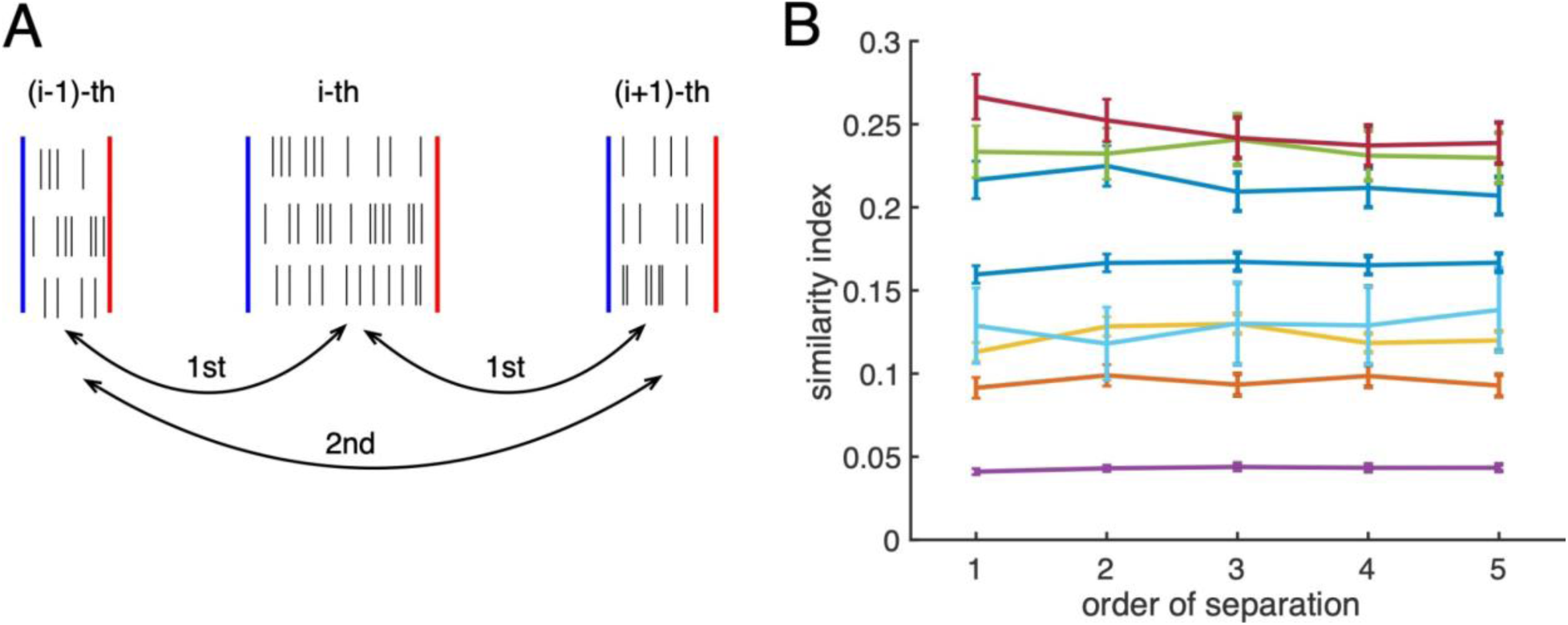
We investigated the temporal diversification of activation patterns, evaluating the similarity index between bursts with 1st – 5th order of separation (schematic illustration shown in panel A). Distributions of similarity index for n=8 samples in the baseline condition suggested that the diversification is temporally independent (B). Note that the tendency was the same for samples under DA conditions (data not shown).

**Supplementary Figure S5:**
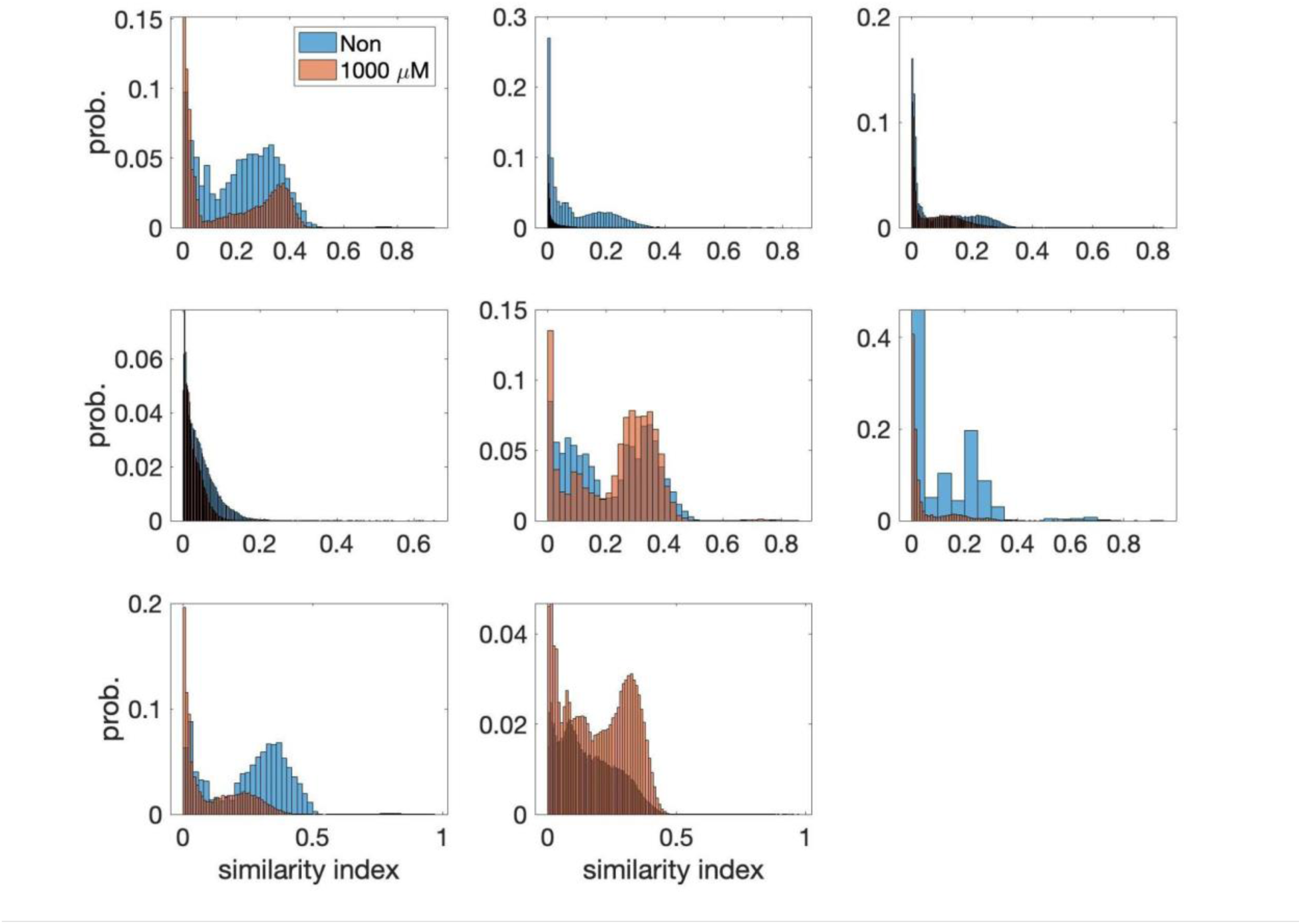
Distributions of similarity index for the 8 samples in the baseline (blue bars) and 1000 μM DA condition (orange bars).

